# Description, Development and Dissemination of Two Consistent Marker-based and Markerless Multibody Models

**DOI:** 10.1101/2022.11.08.515577

**Authors:** Bhrigu Kumar Lahkar, Anaïs Chaumeil, Raphaël Dumas, Antoine Muller, Thomas Robert

**Affiliations:** Univ Lyon, Univ Eiffel, Univ Claude Bernard Lyon 1, LBMC UMR_T9406, F-69622 Lyon, France

**Keywords:** multibody kinematic model, body segment inertial parameters, Theia3D, Visual3D

## Abstract

In human movement analysis, multibody models are an indispensable part of the process both for marker-based and video-based markerless approaches. Constituents (segments, joint constraints, body segment inertial parameters etc.) of such models and modeler’s choice play an important role in the accuracy of estimated results (segmental and joint kinematics, segmental and whole-body center of mass positions etc.). For marker-based method, although standard models exist, particularly for the lower extremity (e.g., Conventional Gait Model or models embedded in OpenSim), there seems to be a lack of consolidated explanation on the constituents of the whole-body model. For the markerless approach, multibody kinematic models (e.g., the Theia3D model) have been in use lately. However, there is no clear explanation on the estimated quantities (e.g., joint centers, body surface landmarks etc.) and their relation to the underlying anatomy. This also motivates the need for a description of the markerless multibody model. Moreover, comparing markerless results to those of classical marker-based method is currently the most commonly used approach for evaluation of markerless approaches. This study first aims to develop and describe a whole-body marker-based model ready to be used for human movement analysis. Second, the markerless multibody model embedded in Theia3D is described and inertial parameters are redefined. We also report assessment of the markerless approach compared to marker-based method for a static T-pose performed by 15 subjects. Finally, we disseminate the marker-based and markerless multibody models for their use in Visual3D.

## 1 Introduction

Human movement is usually quantified through the use of inverse approaches based on human models and tracking external data such as positions of skin markers or velocities and accelerations of inertial sensors. Indeed, direct approaches would require invasive measures such as intracortical pins or medical imaging techniques that remain limited in use (small field of view, non-ecological situations, potentially invasive or irradiant techniques). Kinematics variables are often assessed through the use of multibody kinematic optimization (MKO) (Begon et al., 2018), relying on an a-priori kinematic model of the studied person. This approach appears to be more robust against measurement errors than a classical inverse kinematics approach (i.e. without joint and/or rigid body constraints). However, results are strongly dependent on the kinematic model used (segments considered, constraints used to represent the joints, etc.) (Duprey et al., 2010), and the method used to personalize and to drive the model (i.e., anatomical characteristics of interest and tracked external data). Similarly, internal loads are classically assessed by an inverse dynamic procedure, whose accuracy depends on the correct estimation of the personalized body segment inertial parameters (BSIP) expressed in local segment coordinate systems (SCS), the segments’ kinematics and the position of joint centers when they are used as point of expression for the inter-segmental moments (Camomilla et al., 2017; Derrick et al., 2020).

Standards have emerged for marker-based motion capture approaches. For example, the International Society of Biomechanics (ISB) has provided recommendations regarding the expression of kinematic variables (Wu et al., 2005, 2002). There exist well defined pipelines including a kinematic model, a markerset and the associated processing tools for the lower limb segments (e.g. Conventional Gait Model). Models embedded in freely available tools such as OpenSim have become standards in the biomechanics community (Delp et al., 2007). Still, these standards cover only a part of the process (e.g. ISB recommendations), and are specific to some anatomical region (e.g. lower limbs for the Conventional Gait Model) or are software specific (e.g. OpenSim). As such, many studies rely on alternative “home-made” models. When considering such models, all the necessary characteristics (model parameters and personalization techniques) are often barely described. Indeed, it represents multitude of technical information while it is rarely the main focus of the studies. Therefore, even for these classical marker-based motion capture approaches, there is still need for a whole-body model associated to a clear and citable description of its structure, parameters, and links with the anatomy and with the external markers.

Moreover, with the advent of computer vision and machine learning, new motion capture techniques have come to light, based on calibrated multiview video without markers, later referred to as “markerless”. These are currently based on the detection of the 2D positions of “points of interest” in video via deep-learning algorithms, further transformed in 3D coordinates by triangulation of the 2D views. The 3D trajectories are used to drive a kinematic model through MKO. In most proposed approaches, the link between measured quantities (“points of interest”) and the anatomy is not clearly described (Cao et al., 2019) nor the model used for the inverse kinematic (IK) approaches (Kanko et al., 2021). Here again, there is a need for a well described model, and its association with the underlying anatomy and with the measured quantities. Moreover, it would be particularly interesting if both “marker-based” and “markerless” models could provide comparable outputs. Indeed, markerless approaches are still in a development phase that requires evaluation by comparing their outputs versus classical marker-based results. It would also be essential that results from studies obtained via markerless approaches could be compared to previous results and databases constructed using marker-based approaches.

Therefore, this study aims at providing two whole body models (with male and female versions for each), and their full descriptions, ready to be used for whole-body human movement analysis based either on marker-based or markerless approaches. We also provide insights into the comparison of results obtained from both models.

The markerless approach used is Theia3D (Theia Markerless Inc., Kingston, Ontario, Canada, v2021.2), a commercial human pose estimation software that has recently attained substantial popularity in the field of biomechanics. Models are proposed for use in Visual3D (C-motion, Germantown, USA, v2021.11.3). The following section details the kinematic analysis pipeline of Visual3D and its use of models for both marker-based and markerless approaches.

## 2 Overview of Visual3D kinematic pipeline

Visual3D is a commercial software for calculating kinematic and kinetic variables for biomechanical analysis of 3D motion capture data. **Figure 1** displays the classical pipeline used for kinematic analysis in Visual3D. A subject-specific model is generated from data of a calibration trial (e.g. a static trial) and a model template. Based on the movement trial data (stored in a *c3d* file) and using this subject-specific model, kinematic data (such as joint coordinates or joint positions) are computed by using IK step. Depending on the motion capture approach (marker-based or markerless), the movement data may differ and the model generation and IK steps also differ.

**Figure 1.**
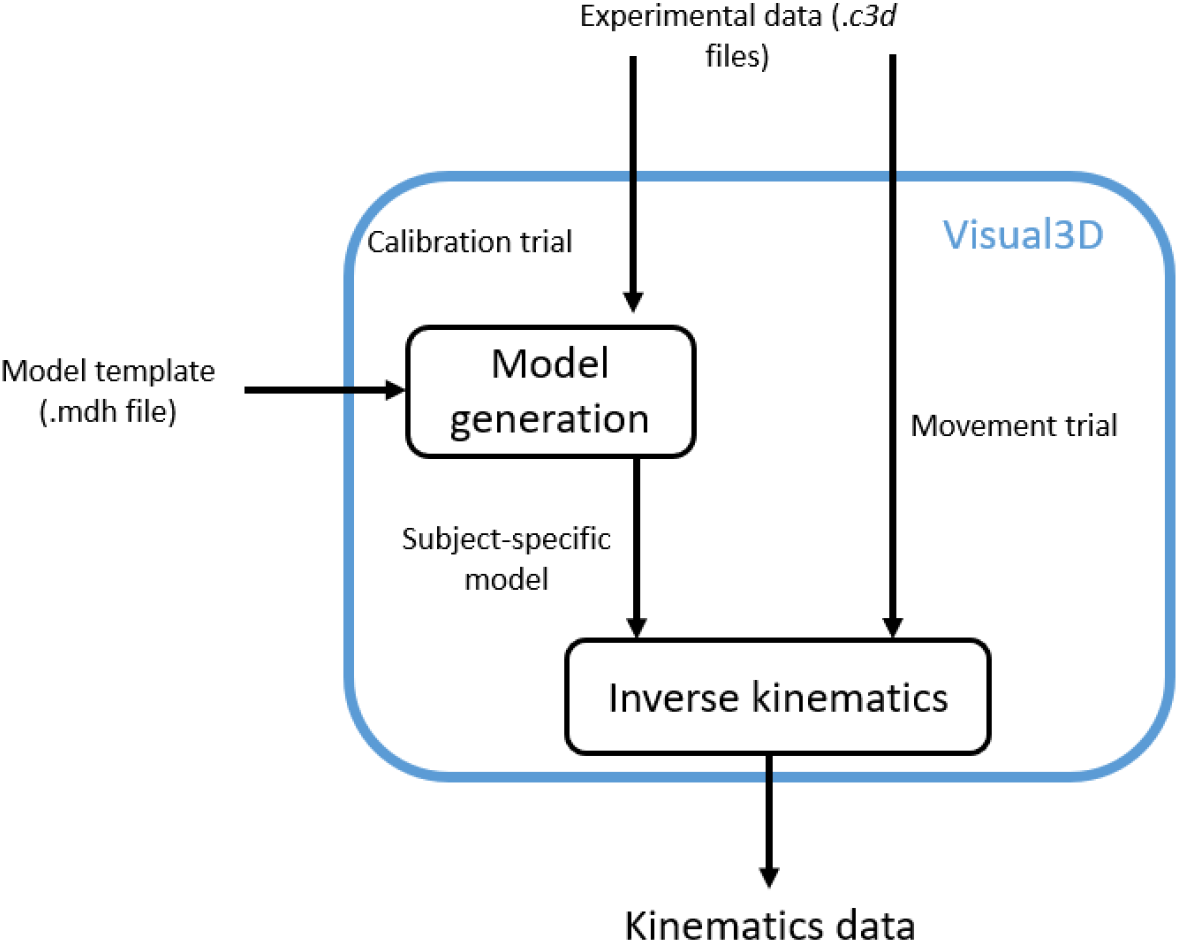
Kinematics analysis pipeline in Visual3D

For the marker-based approach, *c3d* files contain the 3D trajectories of skin markers. Classically by using a static trial, SCS are constructed from the position of anatomical landmarks. BSIP (calculated by means of regression equations), location of model markers and joint constraints are defined to constitute the human model. The IK step consists of an MKO where the model markers are driven by their experimental measured positions over time.

For the markerless approach, *c3d* files are exported fromTheia3D containing pose matrices of each SCS (4×4 matrices containing the position of its origin and the rotation matrix of its base). These segments correspond to a multibody kinematic model predefined in Theia3D which is calibrated and driven by the 3D trajectories of “points of interest”. In Visual3D, SCS are constructed from pose matrices and BSIP are estimated using regression equations. The IK step in Visual3D consists of imposing segment orientation as contained in pose matrices.

For both marker-based and markerless approaches, the model generation is based on a model template file (.*mdh*), containing information on how the model is constructed from the calibration file (.*c3d*). These are the template files that are provided with this paper. A model template is a file that can be opened with any text editor. It contains several sections:

- metrics: constants used to construct the model (participant’s mass and height, segment lengths, ratio for visual, etc.). The participant’s mass is used for BSIP calculation and must be adapted for each participant;
- landmarks: points used to construct the model;
- segments definition: for each segment, SCS construction, BSIP (mass, position of center of mass, inertia), drivers (markers for marker-based or rotation matrix for markerless approach), visual parameters;
- degrees of freedom (only for the marker-based approach): joint constraints used in the MKO;
- optimization algorithm (only for the marker-based approach): parameters for the MKO algorithm.

In Visual3D, SCS is constructed on the basis of 3 landmarks. These landmarks are proximal and distal joint centers of the segment and one extra landmark to define orientation of the segment. The joint centers are synonymously referred to as segment end points. The distal and proximal ends of the segment can be defined explicitly or by providing information of the medial and lateral landmarks. The later method automatically defines a segment end point at the middle of the medial and lateral landmarks.

## 3 Marker-based model

Two marker-based model templates are implemented (one for male ‘ModelMaleMarkerBased.mdh’ and one for female ‘ModelFemaleMarkerBased.mdh’). The models follow the methodology adopted from (Dumas and Wojtusch, 2018). This section describes the marker-set and its anatomical locations followed by descriptions of the model: topology, SCS definition, degree of freedom (DoF), BSIP, and finally the parameters for the IK step.

### 3.1 Marker-set

The marker-set is illustrated in ***Figure 2***, with description of markers in **Table 1**. Technical markers are not used for the model generation but are used as drivers in the MKO. Marker landmarks and its’ nomenclatures are adopted from Color Atlas of Skeletal Landmark Definitions (Van Sint Jan et al., 2007).

**Figure 2.**
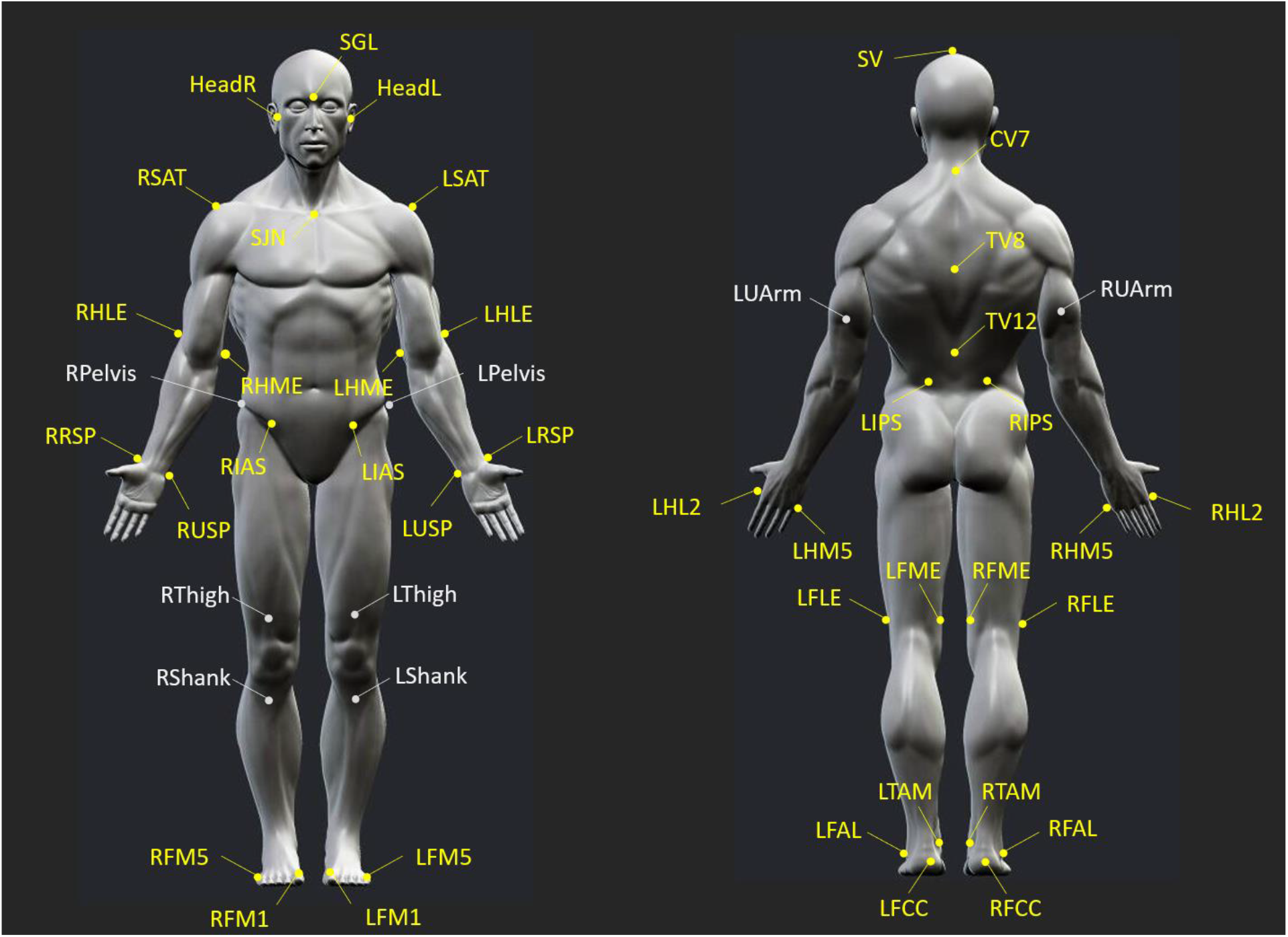
Scheme depicting locations of 48 markers onto the whole-body

**Table 1.** Marker-set description. Markers with * depict technical markers

| **Body Region** | **Acronym** | **Landmark** |
| --- | --- | --- |
| **Head** | SV | Skull Vertex |
|  | SGL | Glabella |
|  | HeadR*, HeadL* | (R, L) Head, at base of the ears |
| **Thorax** | CV7 | 7<sup>th</sup> Cervical vertebra |
|  | TV8 | 8<sup>th</sup> Thoracic vertebra |
|  | TV12 | 12<sup>th</sup> Thoracic vertebra |
|  | SJN | Jugular Notch |
| **Pelvis** | RIAS, LIAS | (R, L) Anterior Superior Iliac Spine |
|  | RIPS, LIPS | (R, L) Posterior Superior Iliac Spines |
|  | RPelvis*, LPelvis* | (R, L) Lateral Superior Iliac Spines |
| **Upper-arm** | RSAT, LSAT | (R, L) Scapula-Acromial Tip |
|  | RHLE, LHLE | (R, L) Humerus Lateral Epicondyle |
|  | RUarm*, Luarm* | (R, L) Upper arm |
|  | RHME, LMLE | (R, L) Humerus Medial Epicondyle |
| **Forearm** | RRSP, LRSP | (R, L) Radius Styloid Process |
|  | RUSP, LUSP | (R, L) Ulna Styloid Process |
| **Hand** | RHL2, LHL2 | (R, L) 2<sup>nd</sup> Metacarpal Head Lateral |
|  | RHM5, LHM5 | (R, L) 5<sup>th</sup> Metacarpal Head Medial |
| **Thigh** | RFLE, LFLE | (R, L) Femoral Lateral Epicondyle |
|  | RFME, LFME | (R, L) Femoral Medial Epicondyle |
|  | RThigh*, LThigh* | (R, L) Thigh |
| **Shank** | RFAL, LFAL | (R, L) Apex of Lateral Malleolus |
|  | RTAM, LTAM | (R, L) Apex of Medial Malleolus |
|  | RShank*, LShank* | (R, L) Shank |
| **Foot** | RFCC, LFCC | (R, L) Foot Calcaneus Center |
|  | RFM1, LFM1 | (R, L) 1<sup>st</sup> Foot Metatarsus |
|  | RFM5, LFM5 | (R, L) 5<sup>th</sup> Foot Metatarsus |

### 3.2 Topology

The marker-based model is composed of 17 segments: head, thorax, clavicles, upper arms, forearms, hands, pelvis, shanks, and feet, separated into two kinematic chains (upper and lower extremity) and a separate head segment (**Figure 3**). This means that free joints (i.e., 6 DoFs) are considered between the global reference system and the pelvis, the global reference system and the thorax, and the global reference system and the head.

**Figure 3.**
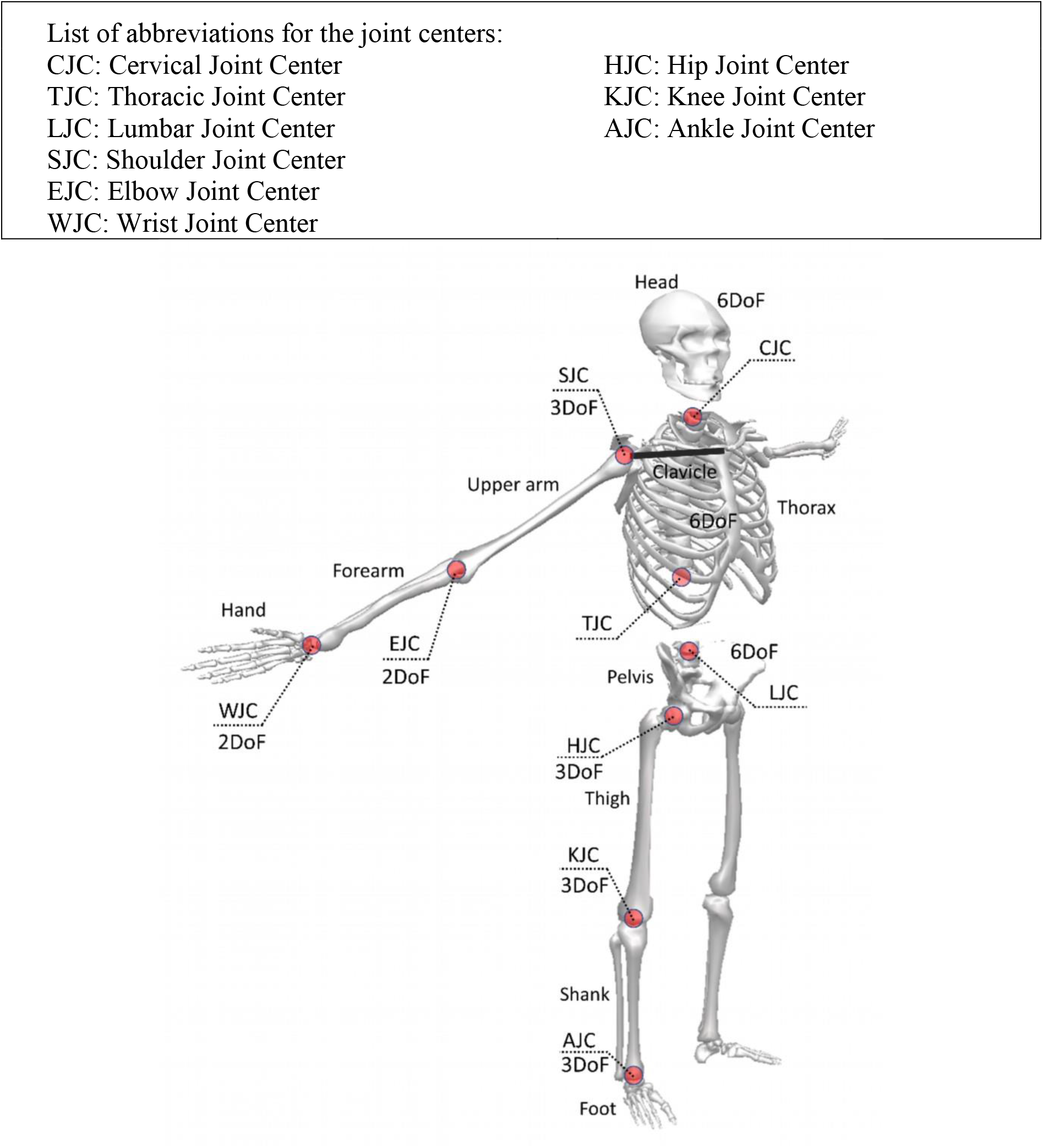
Schematic representation of the multibody model consisting of 2 kinematic chains (upper and lower extremity).

### 3.3 SCS definitions

Here we pictorially detail construction of the SCS from the markers. Construction of the head segment is described in **Figure 4**. Construction of the upper extremity segments (thorax, upper arm, forearm and hand) are described in **Figure 5**. Construction of the lower extremity segments (pelvis, thigh, shank and foot) are described in **Figure 6**.

**Figure 4.**
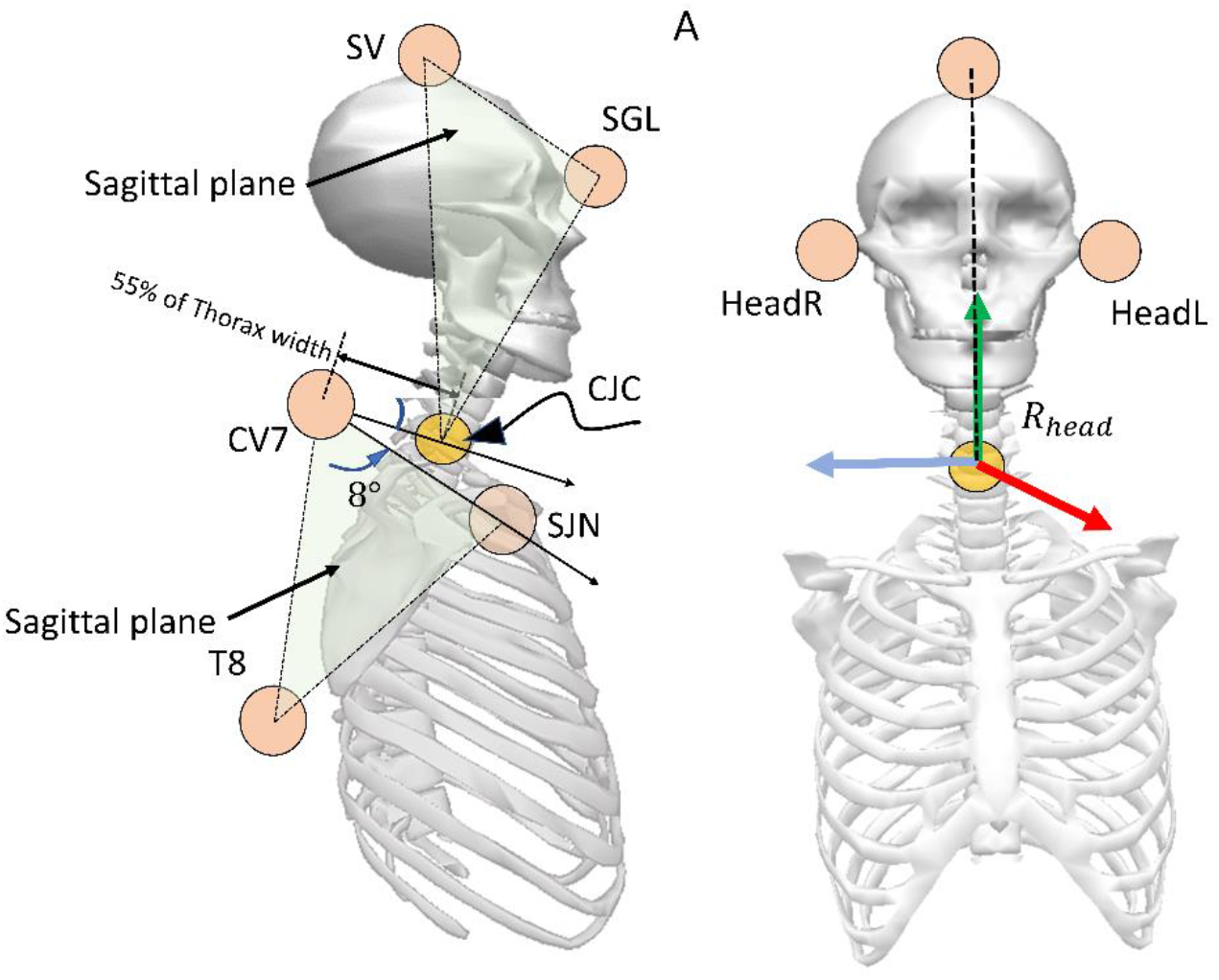
Scheme of the head segment defined between SV and CJC, with origin of segment coordinate system at the CJC. The CJC is defined in a direction forming an angle 𝜽_𝒛_ (male: 8°, female: 14°) counter-clock-wise in the sagittal plane with the vector from CV7 to SJN landmark, and at 𝒙% (male: 55%, female: 53%) of thorax width (distance between CV7 to SJN) from CV7 landmark.

**Figure 5.**
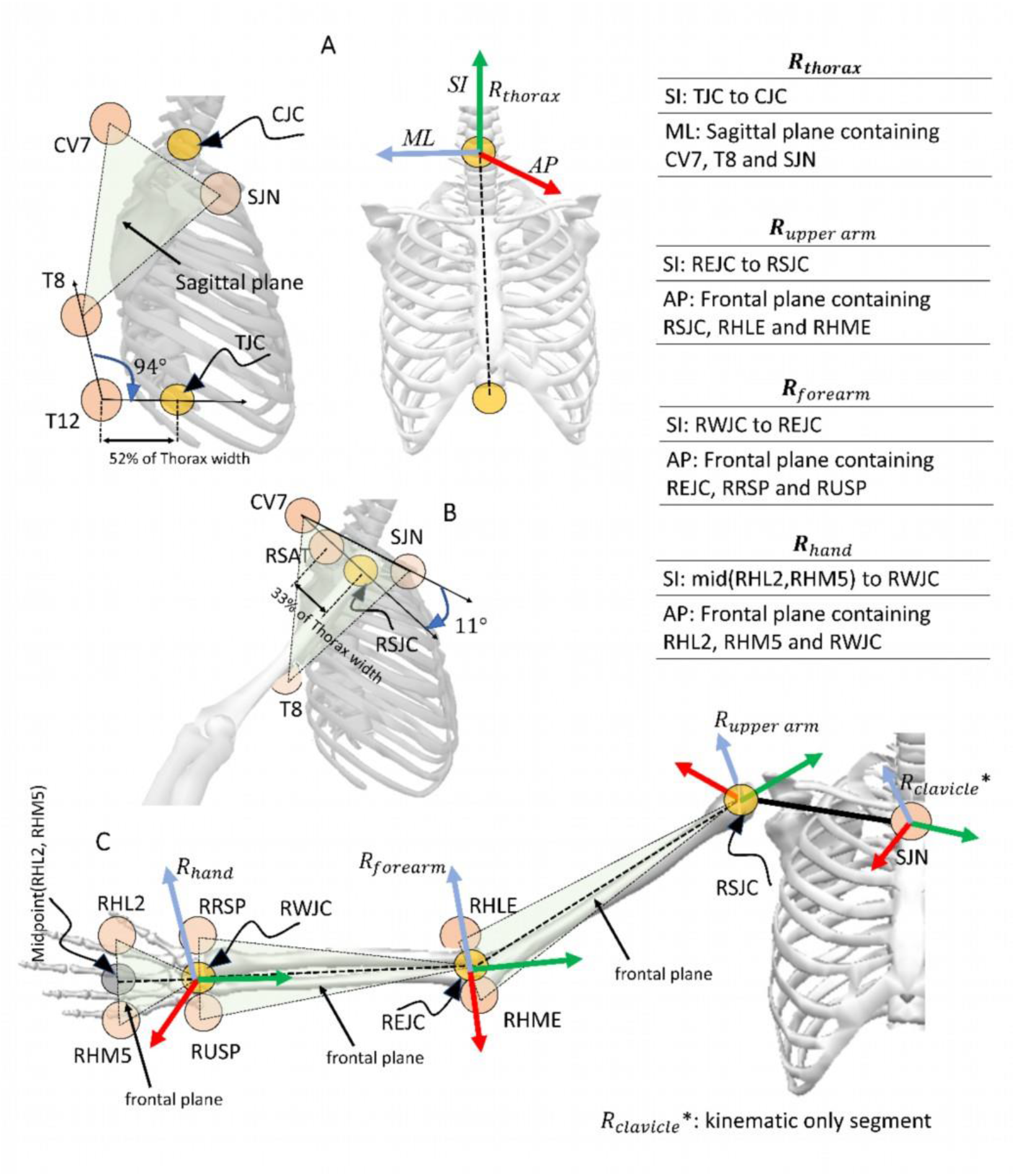
Illustration of the upper extremity segments, their coordinate systems and joint centers **(A)** Thorax segment is defined between TJC and CJC, with origin of the segment coordinate system (*R*_*thorax*_) at the CJC. For definition of CJC, refer to head segment. TJC is defined in a direction forming an angle *θ*_*z*_ (male: 94°, female: 92°) clock-wise in the sagittal plane with the vector from T12 to T8 landmarks, and at *x*% (male: 52%, female: 50%) of thorax width from T12 landmark. **(B)** RSJC is in a direction forming an angle *θ*_*z*_ (male: 11°, female: 5°) clock-wise in the sagittal plane with the vector from CV7 to SJN landmarks, and at *x*% (male: 33%, female: 36%) of thorax width from acromion (RSAT) landmark. Shown for the right shoulder joint. **(C)** The right clavicle is defined between SJN and RSJC, with origin of the segment coordinate system (*R*_*clavicle*_*: kinematic only segment) at the SJN. The right upper arm is defined between RSJC and REJC (midpoint between RHLE and RHME) landmark, with origin of the segment coordinate system (*R*_*upper arm*_) at the RSJC. The right forearm is defined between REJC and RWJC (midpoint between RRSP and RUSP), with origin of the segment coordinate system (*R*_*forearm*_) at REJC. Similarly, the right hand is defined between RWJC and midpoint of RHL2 & RHM5, with origin of the segment coordinate system *R*_*hand*_ at the RWJC. Only shown for right-side joints and segments of the upper extremity.

**Figure 6.**
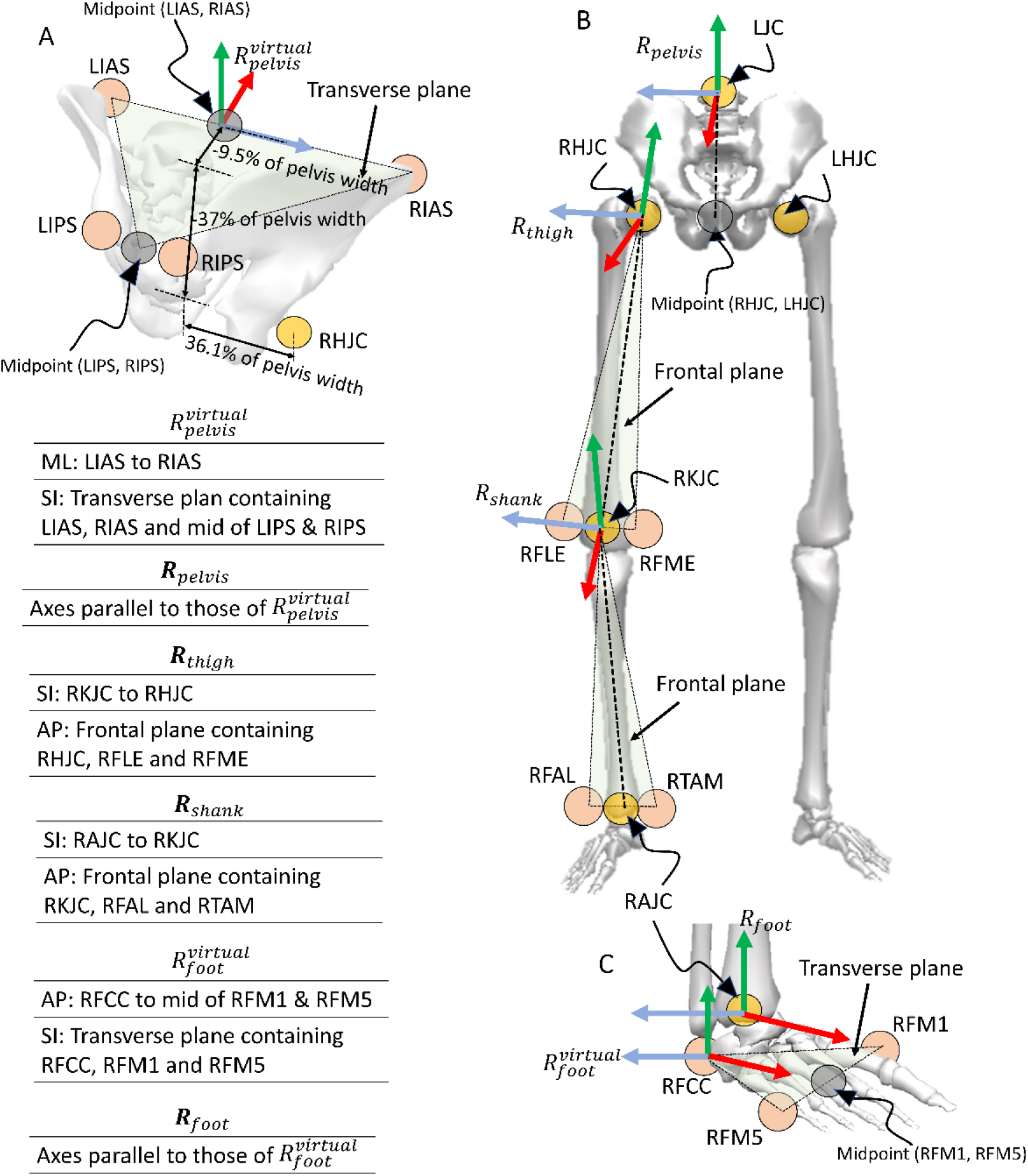
Illustration of the lower extremity segments, their coordinate systems and joint centers **(A)** The right and left hip joint centers (RHJC and LHJC) are defined using regression as: at *x*% (male: −9.5%, female: −13.9%), *y*% (male: −37.0%, female: −33.6%) and at *z*% (male: +/−36.1% (right/left), female: +/−37.2% (right/left)) of the pelvis width (distance between LIAS and RIAS) along AP, SI and ML respectively from the mid-point of LIAS and RIAS landmarks in the Virtual pelvis 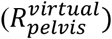. **(B)** The pelvis segment is defined between LJC and mid-point of RHJC & LHJC, with segment axes (*R*_*pelvis*_) parallel to 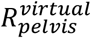 and origin at the LJC. The Right thigh is defined between RHJC and RKJC, with origin of the segment coordinate system (*R*_*thigh*_) at the RHJC. Similarly, the right shank is defined between RKJC and RAJC, with origin of the segment coordinate system (*R*_*shank*_) at the RKJC. **(C)** The right foot is defined between RAJC and the midpoint of RFM1 & RFM5 with origin at the RAJC, and segment axes parallel to the virtual foot 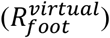. Only shown for the right-side segments of the lower extremity.

#### Construction of the head segment

#### Construction of the upper body segments

#### Construction of the lower body segments

While constructing the body segments, virtual coordinate systems are used for the pelvis 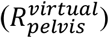 and the feet 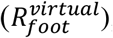, on the basis of which their actual SCS *R*_*pelvis*_ and *R*_*foot*_ have been defined. In the case of the pelvis (*R*_*pelvis*_), it is defined that the mediolateral axis goes “from left to right anterior superior iliac spine landmarks”, with the “transverse plane containing skin landmarks on left and right anterior-superior iliac spines and midpoint between posterior-superior iliac spines”, and with “origin at the lumbar joint center” (Dumas and Wojtusch, 2018). As Visual3D doesn’t directly allow to define such pelvis SCS with origin at lumbar joint center, a virtual SCS is being defined with origin located at the midpoint of the anterior superior iliac spines. The actual pelvis axes are parallel to the respective virtual axes, but the origin is defined at the lumbar joint center. The process is similar for the feet on each side. For each foot, as described in (Dumas and Wojtusch, 2018), the “anterior-posterior axis goes from heel to midpoint between 1^st^ and 5^th^ metatarsal head skin markers” with origin at the AJC. It is not possible to define such foot SCS directly in Visual3D, thus a virtual foot is defined with origin at heel marker and anterior-posterior axis from heel to midpoint between metatarsal head markers. The actual foot axes are parallel to the virtual axes, but with origin defined at the AJC.

### 3.4 Joint DoFs

The DoFs assigned for each joint are defined in **Table 2**. The rotation sequences are based on ISB recommendations (Wu et al., 2005, 2002) to be consistent with the definition of the model.

**Table 2:** DoFs for each joint.

| Joint (segment relative to parent segment) | Rotation sequence |
| --- | --- |
| Hip (thigh relative to the pelvis) | ZX’Y’’ |
| Knee (shank relative to the thigh) | ZX’Y’’ |
| Ankle (foot relative to the shank) | ZY’X’’ |
| Shoulder 1* (clavicle relative to the thorax) | YX’ |
| Shoulder 2 (upper arm relative to the clavicle) | YX’Y’’ |
| Elbow** (forearm relative to the upper arm) | ZY’’ |
| Wrist (hand relative to the forearm) | ZX’’ |

### 3.5 BSIP

The inertial parameters for all the segments: segment mass, position of center of mass (CoM), inertia are adopted from (Dumas and Wojtusch, 2018), and are detailed in **Table 3** and **Table 4**.

**Table 3:** Segment properties of the marker-based upper extremity model including the head

| Segment name | Segment ends for segment length calculations<br>Proximal, Distal | Segment tracking markers | Segment mass<br>(% of total body mass)<br>(female, male) | Position of CoM<br>(% of segment length)<br>(female, male) | Moment of inertia<br>(radius of gyration in % of segment length)<br>(female, male) |
| --- | --- | --- | --- | --- | --- |
| Head | CJC<br>SV | HeadR,<br>HeadL,<br>SGL, SV | 6.7, 6.7 | AP: 0.8, 2.0 | AP: 30, 28 |
|  |  |  |  | SI: 55.9, 53.4 | SI: 24, 21 |
|  |  |  |  | ML: -0.1, 0.1 | ML: 31, 30 |
| Thorax | CJC<br>LJC | SJN, CV7,<br>TV8, TV12 | 30.4,<br>33.3 | AP: -1.6, -3.6 | AP: 29, 27 |
|  |  |  |  | SI: -43.6, -42.0 | SI: 27, 25 |
|  |  |  |  | ML: -0.6, -0.2 | ML: 29, 28 |
| Clavicle<br>(kinematic only) | SJN<br>SJC | SJN, SAT | - | - | - |
| Upper arm | SJC<br>EJC | HLE,<br>HME,<br>UArm | 2.3, 2.4 | AP: -5.5, 1.8 | AP: 30, 29 |
|  |  |  |  | SI: -50.0, -48.2 | SI: 15, 13 |
|  |  |  |  | ML: -3.3, -3.1 | ML: 30, 30 |
| Forearm | EJC<br>WJC | USP, RSP | 1.4, 1.7 | AP: 2.1, -1.3 | AP: 27, 28 |
|  |  |  |  | SI: -41.1, -41.7 | SI: 14, 11 |
|  |  |  |  | ML: 1.9, 1.1 | ML: 25, 28 |
| Hand | WJC<br>Mid (HL2, HM5) | HL2, HM5 | 0.5, 0.6 | AP: 7.7, 8.2 | AP: 64, 61 |
|  |  |  |  | SI: -76.8, -83.9 | SI: 43, 38 |
|  |  |  |  | ML: 4.8, 7.5 | ML: 59, 56 |

**Table 4:** Segment properties of the marker-based lower extremity model. * For the foot in Visual3D, the axial direction corresponds to the anatomical AP direction. Thus, values corresponding to anatomical AP direction are associated with Visual3D axial direction, and values corresponding to anatomical SI direction are associated to Visual3D AP direction. Only the values along ML direction remain unchanged.

| Segment name | Segment ends for segment length calculations<br>Proximal, Distal | Segment tracking markers | Segment mass (% of total body mass)<br>(female, male) | Position of CoM (% of segment length)<br>(female, male) | Moment of inertia (radius of gyration in % of segment length)<br>(female, male) |
| --- | --- | --- | --- | --- | --- |
| Pelvis | LJC<br>Mid (RHJC, LHJC) | RIAS,<br>LIAS,<br>RIPS, LIPS | 14.7, 14.2 | AP: -7.2, -0.2 | AP: 95, 102 |
|  |  |  |  | SI: -22.8, -28.2 | SI: 105, 106 |
|  |  |  |  | ML: 0.2, -0.6 | ML: 82, 96 |
| Thigh | HJC<br>KJC | FLE, FME,<br>Thigh | 14.6, 12.3 | AP: -7.7, -4.1 | AP: 31, 29 |
|  |  |  |  | SI: -37.7, -42.9 | SI: 19, 15 |
|  |  |  |  | ML: 0.8, 3.3 | ML: 32, 30 |
| Shank | KJC<br>AJC | FAL,<br>TAM,<br>Shank | 4.5, 4.8 | AP: -4.9, -4.8 | AP: 28, 28 |
|  |  |  |  | SI: -40.4, -41.0 | SI: 10, 10 |
|  |  |  |  | ML: 3.1, 0.7 | ML: 28, 28 |
| Foot | AJC<br>Mid (FM1, FM5) | FCC, FM1,<br>FM5 | 1.0, 1.2 | AP*: 38.2, 50.2 | AP*: 24, 22 |
|  |  |  |  | SI*: -30.9, -19.9 | SI*: 50, 49 |
|  |  |  |  | ML: 5.5, 3.4 | ML: 50, 48 |

Positions of CoM and components of the inertia matrix are calculated by using the segment lengths which are defined by means of segment endpoints. However, there is an exception for the thorax: the distal endpoint (TJC) is not used for segment length calculation. This is due to the absence of a separate abdomen segment as proposed in (Dumas and Wojtusch, 2018), which led us to define the thorax segment length from CJC to LJC, as the torso defined in (Dumas et al., 2007). The position of CoM and components of the inertia matrix are expressed using this and BSIP defined in (Dumas et al., 2007). Mass of the torso is also expressed using (Dumas et al., 2007).

### 3.6 IK parameters

Tracking markers are associated to segments and are used as drivers during the IK step (MKO). Information about tracking markers are provided in **Table 3** and **Table 4**.

We also provide segment-specific marker weights based on residual analysis for the movement under study. These weights can be found directly in the mdh files. The users are encouraged to adjust them according to their study. The global optimization algorithm is selected as Quasi-Newton by default.

## 4 Markerless model

Two markerless model templates are implemented (one for male ‘ModelMaleMarkerLess.mdh’ and one for female ‘ModelFemaleMarkerLess.mdh’). Their topology and SCS definition are those of the model of Theia3D (https://www.theiamarkerless.ca/docs/model.html), and BSIP adapted from (Dumas and Wojtusch, 2018)

### 4.1 Topology

The markerless model is composed of 17 segments: head, thorax, upper arms, forearms, hands, pelvis, thighs, shanks, feet and toes, separated into two kinematic chains (upper and lower extremity). This means that free joints (6 DoFs) are considered between the global reference system and the pelvis, the global reference system and the thorax, and the global reference system and the head.

### 4.2 SCS definitions

SCS are constructed based on the pose matrices. **Table 5** and **Table 6** describe how the different SCS are built.

**Table 5:** Segment properties of the markerless upper extremity model including the head

| Segment name | Segment ends<br>Proximal:<br>Distal: | Segment mass<br>(% of total<br>body mass)<br>(female, male) | Position of CoM<br>(% of segment<br>length)<br>(female, male) | Moment of inertia<br>(radius of gyration in<br>% of segment length)<br>(female, male) |
| --- | --- | --- | --- | --- |
| Head | RHE_Proximal<br>Head_4x4 | 6.7, 6.7 | AP: 0.8, 2.0 | AP: 30, 28 |
|  |  |  | SI: 55.9, 53.4 | SI: 24, 21 |
|  |  |  | ML: -0.1, 0.1 | ML: 31, 30 |
| Thorax | Torso_4x4<br>RTX_distal | 30.4<br>33.3 | AP: -1.6, -3.6 | AP: 29, 27 |
|  |  |  | SI: -43.6, -42.0 | SI: 27, 25 |
|  |  |  | ML: -0.6, -0.2 | ML: 29, 28 |
| Upper arm | R_uarm_4x4<br>R_larm_4x4 | 2.3, 2.4 | AP: -5.5, 1.8 | AP: 30, 29 |
|  |  |  | SI: -50.0, -48.2 | SI: 15, 13 |
|  |  |  | ML: -3.3, -3.1 | ML: 30, 30 |
| Forearm | R_larm_4x4<br>R_hand_4x4 | 1.4, 1.7 | AP: 2.1, -1.3 | AP: 27, 28 |
|  |  |  | SI: -41.1, -41.7 | SI: 14, 11 |
|  |  |  | ML: 1.9, 1.1 | ML: 25, 28 |
| Hand | R_hand_4x4<br>RHA_Distal | 0.5, 0.6 | AP: 7.7, 8.2 | AP: 64, 61 |
|  |  |  | SI: -76.8, -83.9 | SI: 43, 38 |
|  |  |  | ML: 4.8, 7.5 | ML: 59, 56 |

**Table 6:** Segment properties of the markerless lower extremity model. * For the foot in Visual3D, the axial direction corresponds to the anatomical AP direction. Thus, values corresponding to anatomical AP direction are associated with Visual3D axial direction, and values corresponding to anatomical SI direction are associated to Visual3D AP direction. Only the values along ML direction remain unchanged.

| Segment name | Segment ends<br>Proximal:<br>Distal: | Segment mass<br>(% of total<br>body mass)<br>(female, male) | Position of CoM<br>(% of segment<br>length)<br>(female, male) | Moment of inertia<br>(radius of gyration in<br>% of segment length)<br>(female, male) |
| --- | --- | --- | --- | --- |
| Pelvis | Pelvis_shifted_4x4<br>Mid_hip | 14.7, 14.2 | AP: -7.2, -0.2 | AP: 95, 102 |
|  |  |  | SI: -22.8, -28.2 | SI: 105, 106 |
|  |  |  | ML: 0.2, -0.6 | ML: 82, 96 |
| Thigh | Rthigh_4x4<br>Rshank_4x4 | 14.6, 12.3 | AP: -7.7, -4.1 | AP: 31, 29 |
|  |  |  | SI: -37.7, -42.9 | SI: 19, 15 |
|  |  |  | ML: 0.8, 3.3 | ML: 32, 30 |
| Shank | R_shank_4x4<br>R_foot_4x4 | 4.5, 4.8 | AP: -4.9, -4.8 | AP: 28, 28 |
|  |  |  | SI: -40.4, -41.0 | SI: 10, 10 |
|  |  |  | ML: 3.1, 0.7 | ML: 28, 28 |
| Foot | R_foot_4x4<br>RFT_Distal | 1.0, 1.2 | AP*: 38.2, 50.2 | AP*: 24, 22 |
|  |  |  | SI*: -30.9, -19.9 | SI*: 50, 49 |
|  |  |  | ML: 5.5, 3.4 | ML: 50, 48 |
| Toes | R_toes_4x4<br>Rtoes_distal | 10<sup>-8</sup>, 10<sup>-8</sup> | AP: 10<sup>-8</sup>, 10<sup>-8</sup> | AP: 0, 0 |
|  |  |  | ML: 10<sup>-8</sup>, 10<sup>-8</sup> | ML: 0, 0 |
|  |  |  | SI: 10<sup>-8</sup>, 10<sup>-8</sup> | SI: 0, 0 |

### 4.3 DoFs

The following DoFs are considered in the Theia3D kinematic model: 3 DoFs for the hip (thigh relative to the pelvis), 3 DoFs for the knee (shank relative to the thigh), 3 DoFs for the ankle (foot relative to the shank), 1 DoF for metatarsophalangeal joint (toe relative to the foot) (flexion/extension), 2 DoFs for the shoulder1 (clavicle relative to the thorax) (about anterior-posterior and superior-inferior axes), 3 DoFs for the shoulder2 (upper arm relative to the clavicle), 2 DoFs for the elbow (forearm relative to the upper am) (flexion/extension, pronation/supination), and 2 DoFs for the wrist (hand relative to the forearm) (flexion/extension, ad/abduction). Alternatives to the number of DoFs for some joints (knee and shoulder) exist in Theia3D.

### 4.4 BSIP

The BSIP for all the segments: segment mass, position of CoM and moment of inertia are detailed in **Table 5** and **Table 6**. These values are adopted from (Dumas and Wojtusch, 2018). We did our best to identify how SCS were constructed based on Theia3D documentation and to adapt the BSIP definition accordingly.

Positions of CoM and components of the inertia matrix are calculated by using the segment lengths. which are defined by means of segment endpoints. For thighs, shanks, upper arms, and forearms, the segment length is computed as the distance between the proximal and distal endpoints, based on the data from the pose matrices. For other segments (hands, feet, pelvis, thorax, and head), mostly because of the lack of appropriate distal endpoint, segment length is calculated as the product of a precomputed ratio (eq. 1) and its standard length coming from (Dumas and Wojtusch, 2018).

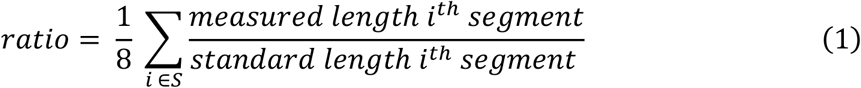

Where *S* includes thighs, shanks, upper arms, and forearms segments.

In (Dumas and Wojtusch, 2018), there is no distinction made between the feet and toes, thus the BSIP of toes are set to zero.

As the abdomen segment is not considered in the model, the BSIP of both thorax and abdomen defined in (Dumas and Wojtusch, 2018) are assigned to the thorax segment. The BSIP are based on (Dumas et al., 2007), for which the thorax is defined between the CJC (“base of neck” in Theia3D model) and LJC (pelvis_shifted_4×4). The length of the thorax is also computed based on this definition.

For the pelvis, the origin of the SCS is set at the LJC. An assumption is made for the markerless model, stating that the origin of the pelvis SCS is at a point corresponding to the LJC.

For the head and thorax, an assumption is made that the CJC defined in (Dumas and Wojtusch, 2018) corresponds to the point at the “base of the neck” pointed by the torso pose matrix. These assumptions allow us to use these points, which are defined literally, as if they were built using markers and regression equations, which are expected in (Dumas and Wojtusch, 2018).

## 5 Comparison between marker-based and markerless models

### 5.1 Principle

In the previous sections we described how marker-based and markerless models are defined. If these models can be used independently, it would be interesting if both models could provide comparable outputs. The aim of this section is to compare marker-based and markerless approaches by comparing results of kinematics analysis (segment rotation and position of their center of mass in the global coordinate system) of a static T-pose for several participants.

Fifteen young and healthy participants (8 men, 7 women, mean age: 25.5±2.7 years old, mean height: 1.75±0.09 m, mean mass: 69.8±12.8 kg) were recruited and performed a static T-pose with the palms of their hands directed towards the ground for a few seconds. Participants signed an informed consent form prior to the experiment.

Two camera systems were set up, synchronized and spatially calibrated: a marker-based optoelectronic system composed of 10 Qualisys Miqus M3 cameras (sampling frequency: 300 Hz) and a markerless system composed of 10 Qualisys Miqus Video cameras (sampling frequency: 60 Hz, cameras resolution: 1920 × 1088 pixels). The participants were equipped with the marker-set described in Section 3.1.

Data from marker-based system were labelled and gap filled in Qualisys Track Manager (QTM – Qualisys AB, Sweden, v2021.1.2). The output *c3d* files were imported in Visual3D. Then, the kinematics analysis pipeline detailed in **Figure 1** was followed using the marker-based model detailed in Section 3. Data from video cameras were processed in Theia3D, and the output *c3d* files were imported in Visual3D. Then, the kinematic analysis pipeline detailed in **Figure 1** was followed using the markerless model detailed in Section 4.

Both models were compared firstly in terms of structure (topology, orientation of the SCS in relation to anatomical directions), and secondly in terms of kinematics results (SCS orientations and CoM positions for each segment according to the global coordinate system).

Concerning kinematics results, rotation matrices and CoM positions were extracted from Visual3D for both approaches. For each segment, we estimated the relative orientation between marker-based and markerless SCS. The rotation matrix of markerless SCS with respect to marker-based SCS was computed and the corresponding angles were extracted using an ZX’Y” Euler angle sequence (Wu and Cavanagh, 1995). Average rotation angles along Z, X’, and Y’’ and their standard deviation were calculated across all subjects. Moreover, signed differences of CoM in all three directions of the global coordinate system (X, Y and Z) were computed, and an average value and standard deviation were provided across all subjects. All segments were considered except clavicles where BSIP are zero, and toes which don’t exist in the marker-based model. All these computations were done in MATLAB (MathWorks, USA).

### 5.2 Comparison of the topologies

Firstly, regarding differences of topology, there are toes in the markerless model but not for the marker-based model. However, toes could be added to the marker-based model to add 1 DoF between the foot and toes.

Also, there is a difference in the orientation of the SCS in relation to anatomical directions for both models (**Table 7**). The marker-based model follows the ISB recommendations while this is not the case for the markerless model. A permutation matrix during the post processing was applied to the SCS of the markerless model to allow a consistent comparison of kinematics data for the following section.

**Table 7:** Orientation of SCS according to anatomical direction for both marker-based and markerless models

|  | Marker-based model | Markerless model |
| --- | --- | --- |
| **Antero-posterior** | X | Y |
| **Medio-lateral** | Z | X |
| **Vertical** | Y | Z |

### 5.3 Relative SCS orientation

**Table 8** and **Table 9** present respectively the results of the comparison of SCS orientation and position of CoM of all segments composing marker-based and markerless models during the static pose.

**Table 8:** Difference (Mean±SD) in orientation between the markerless and marker-based SCS, expressed in degrees

| Segment | Along Z (°) | Along X' (°) | Along Y'' (°) |
| --- | --- | --- | --- |
| Pelvis | -8.5±5.6 | 0.2±1.6 | 0.5±3.4 |
| Thorax | -6.4±3.7 | -0.1±2.0 | 2.1±1.8 |
| Head | -33.5±3.5 | -1.4±2.7 | 2.9±2.4 |
| Right thigh | -3.2±2.3 | -0.5±1.1 | 4.7±6.2 |
| Left thigh | -3.1±2.2 | 0.6±1.7 | 1.2±7.2 |
| Right shank | -0.2±1.6 | -0.7±0.9 | -13.5±12.6 |
| Left shank | -1.0±1.2 | 0.6±0.6 | 15.3±10.7 |
| Right foot | -1.7±2.0 | 7.3±4.3 | 1.6±2.1 |
| Left foot | -2.9±1.8 | -3.8±2.2 | -4.1±1.5 |
| Right upper arm | -11.6±2.4 | -0.7±3.9 | 8.2±17.6 |
| Left upper arm | -9.2±3.7 | -5.1±5.0 | 12.9±22.7 |
| Right forearm | 1.3±3.3 | -6.1±2.5 | 22.8±21.0 |
| Left forearm | 0.4±3.7 | 4.5±2.6 | -28.3±20.8 |
| Right hand | -13.5±6.8 | 4.4±7.5 | 20.8±19.8 |
| Left hand | -13.3±9.8 | -5.0±5.1 | -27.5±19.8 |

**Table 9:** Signed difference (marker-based - markerless) of CoM position (in mm) for each segment expressed as Mean±SD. X is pointing forward, Y is pointing upward and Z is pointing to the right side of the participants.

| Segment | X (mm) | Y (mm) | Z (mm) |
| --- | --- | --- | --- |
| **Pelvis** | -23.6±14.7 | -2.9±13.9 | -0.1±3.2 |
| **Thorax** | -25.2±15.2 | -5.3±13.8 | -1.1±7.4 |
| **Head** | 23.8±7.4 | 7.5±11.2 | -0.1±3.0 |
| **Right thigh** | 27.3±8.8 | -5.8±10.6 | 8.9±5.9 |
| **Left thigh** | 24.7±6.2 | -4.9±11.9 | -7.4±8.1 |
| **Right shank** | 11.9±6.8 | 8.2±3.6 | 12.4±4.2 |
| **Left shank** | 9.8±6.5 | 8.9±3.4 | -10.6±3.4 |
| **Right foot** | 4.5±5.5 | 1.3±5.0 | 10.5±4.0 |
| **Left foot** | 0.5±4.0 | 2.8±4.9 | -10.4±2.5 |
| **Right upper arm** | 8.0±9.9 | 2.6±5.5 | 14.5±7.5 |
| **Left upper arm** | 8.3±9.0 | 3.5±6.4 | -13.1±7.5 |
| **Right forearm** | -2.0±7.4 | -18.5±4.9 | 5.7±6.2 |
| **Left forearm** | -3.0±6.3 | -17.8±6.1 | -7.8±6.7 |
| **Right hand** | 1.7±4.6 | -6.4±4.5 | -5.7±6.7 |
| **Left hand** | -5.4±5.0 | -6.6±3.9 | 6.0±7.3 |
| **Whole-body model** | -1.6±6.7 | -2.8±7.0 | -0.1±2.5 |

Differences appear in orientation of the pelvis and thorax along the Z axis (medio-lateral) between the two models. In the marker-based model, the pelvis frame is defined as follows: the medial-lateral axis is defined “from left to right anterior superior iliac spine skin landmarks”, and the transverse plane is defined to contain “skin landmarks on left and right anterior-superior iliac spines and midpoint between posterior-superior iliac spines” (Dumas and Wojtusch, 2018) This leads to an anterior tilt of the pelvis with respect to the global coordinate system which depends on the pose of participant and the placement of the markers. However, for markerless model, it appears that the orientation of the pelvis is defined such that there is no tilt in standing posture, following Visual3D recommendation (“Segment Examples 5 — Visual3D Wiki Documentation”). It thus leads to a difference of initial pelvic tilt with respect to the global coordinate system of around 8°. The associated standard deviation of around 5° remains small, suggesting that the difference of 8° in tilt could be seen as a constant offset, and not a result of a measurement noise.

Regarding the thorax coordinate system, it appears to be defined using similar endpoints for markerless and marker-based methods, although the definition of the endpoints for the markerless model remains approximative. We found a difference of about 6° for the thorax in static position, with a small standard deviation as well.

When using the markerless model, one should be careful on the differences in pelvis and thorax orientation, because both segments are usually the root segments of the lower or upper limbs kinematic chain: an initial tilt would result in changes of joint angles. However, it is possible to take these differences into account in the post processing phase by adding an offset to correct them.

The head displays large difference in orientation along the Z-axis between the two models, yet this difference seems consistent (small SD of about 3°). To the best of our knowledge, the difference of orientation of the head between marker-based and markerless models has never been studied in the literature concerning Theia3D. Two possibilities can explain this result: the keypoints used to detect and orient the head properly are not enough to provide an accurate result. Another option is that, as the participants wore uniform swim caps for easier marker placement, ears were hidden, which could lead to fewer keypoints detected. However, not knowing the keypoints used by Theia3D, it is not possible to conclude.

Both upper arms and hands also displayed differences of orientation along the Z axis of around 10°. For upper arms, a reasonable explanation would be the difficulty to accurately locate the shoulder joint in the video. Furthermore, the T pose of the participants could have generated inaccuracies due to high humerus elevation. In (Lahkar et al., 2022), shoulder angles (between upper arms and thorax) had RMSD of around 10°, which is consistent with our results.

For hands, it is only possible to assume that Theia3D detects keypoints on the hands, and especially fingers. This assumption being made, needless to say that it is difficult to detect accurately these keypoints as the hands are quite small segments and the neural networks can make confusions between two similar looking keypoints. (Lahkar et al., 2022) also found errors of wrist flexion/extension between 14° and 20°.

Regarding internal/external rotation differences between both models (along Y”), the main differences occur for the shank and upper limb segments. Without additional knowledge of the keypoints used to define the shank segment, no hypothesis explaining this difference can be made at first. (Kanko et al., 2021) found, for rotation around the axial axis, a RMSD of 13° for the shank during gait, which is also consistent with our results. For the upper limb segments, such differences can be explained by the position of the participants: as the arms are stretched, it is difficult to orient correctly the segment frames, even with markers. A static posture with flexed elbows could have provided a better correspondence of internal/external rotation between the two models. (Lahkar et al., 2022) also found large RMSD (between 18° and 23°) for elbow internal/external rotation. It is also noticeable that the standard deviations associated to these segments for internal/external rotation are large, which suggests a great variability in the orientation of these segments when using the markerless model.

### 5.4 Comparison of CoM positions in the global coordinate system

Overall, the signed differences between the whole-body CoM obtained with both models are very small in all three directions. This observation is rather counterintuitive when looking at the segmental CoM position errors in X direction: pelvis, thorax, head and thighs all display errors that are above 20 mm, and shanks display errors of around 10 mm. However, when looking at the percentage of mass represented by these segments with major errors, they represent approximately 43% of total body mass for head, thighs and shanks, and 42% of total body mass for pelvis and thorax, the first group bringing a positive difference, and the second group a negative difference between segment CoM locations. It is thus very likely that these two groups, representing a similar part of the total body mass, compensate each other’s error in computation of the whole-body CoM, resulting in very small errors between markerless and marker-based models. Except for the pelvis and thorax, the standard deviations associated with the considered segments in X direction are in overall small compared to the measured differences. However, the obtained values suggest an offset for the segmental CoM position for the aforementioned segments.

Differences of around 10 mm can be observed for shanks, feet and upper arms along the Z direction (medio-lateral). As the arms are extended and directed on both sides of the body in the static T-pose, the difference in global Z direction actually corresponds to a difference in the longitudinal axis of the upper arm. Regarding shanks and feet, these differences are opposed and thus compensate in the whole-body center of mass computation for this static T-pose.

Along the Y direction, only the forearms display differences close to 20 mm. Similarly, to the upper arm in the previous paragraph, a difference in the Y direction observed for the forearms in the static T-pose actually corresponds to a difference in medio-lateral direction in forearm reference frame. For the pelvis, thorax, head and thighs, standard deviations are larger than the measured differences, suggesting a large variability in estimating segmental CoM positions in Y direction.

## 5. Conclusion

In this study, we described two whole body models that are ready to be used for whole body human movement analysis based either on marker-based or markerless approaches. These models are attached to this paper for a use in Visual3D.

We also provided insight into the comparison of results obtained from both approaches (marker-based and markerless) using these models. It showed an overall good agreement but it also highlighted some specific consistent differences, which may be corrected using offsets, and some more variable differences that could be attributed to the measure and for which analyses should be more careful.

**Guideline and recommendation for using model templates\***

#### Marker-based models

The users of the marker-based model templates (.mdh files) are strictly recommended to use the same marker-set and their nomenclature. In case the users prefer their own marker set and nomenclature, sufficient care must be taken to modify those in the editable .mdh files. Any wrong correspondence of the marker-data with the model template will inevitably fail in executing the process in Visual3D. The tracking markers assigned in the given model can be changed (add or remove) based on the study.

We also provide an example calibration file (Example_Static.c3d) in the supplementary material, for a test execution of the female model template. The users should, however, use their calibration file in their studies.

#### Markerless models

The .c3d files exported from Theia can be opened in Visual3D, where a default multibody model is already loaded. To use the markerless model provided with this article in Visual3D, users should click on *Model > Apply model template* and select the appropriate model template, e.g. male or female.

#### All the models can be downloaded from the supplementary material of this article

*\*Please cite this article if you use the models in your research*

## Supporting information

Multibody models

